# Integrative omics analysis reveals gene regulatory mechanisms distinguishing organoid-derived hepatocytes from primary human hepatocytes

**DOI:** 10.1101/2023.12.05.570132

**Authors:** Haoyu Wu, Annie S. P. Yang, Suzan Stelloo, Floris J. M. Roos, René H. M. te Morsche, Anne H. Verkerk, Maria V. Luna-Velez, Laura Wingens, Johannes H. W. de Wilt, Robert W. Sauerwein, Klaas W. Mulder, Simon J. van Heeringen, Monique M. A. Verstegen, Luc J. W. van der Laan, Hendrik Marks, Richárd Bártfai

## Abstract

**Background and Aims:** Hepatic organoid cultures are considered a powerful model system to study liver development and diseases *in vitro*. However, hepatocyte-like cells differentiated from such organoids remain immature compared to primary human hepatocytes. Therefore, a comprehensive understanding of differences in gene regulatory mechanisms between primary human hepatocytes and hepatic organoids is essential to obtain functional hepatocyte-like cells *in vitro* for fundamental and therapeutic applications.

**Methods:** We obtained primary human hepatocytes at high purity from all zones of the liver lobule using an optimized two-step perfusion protocol. We captured the single-cell transcriptome and chromatin accessibility landscape using scRNA-seq and ATAC-seq, respectively. We identified key transcription factors and compared the gene regulatory mechanisms in primary human hepatocytes and (un)differentiated intrahepatic cholangiocyte organoids. Using siRNA-mediated perturbations, we showed the functional relevance of an organoid-enriched transcription factor during *in vitro* differentiation of hepatocyte-like cells.

**Results:** Our integrative omics analysis revealed that Activator Protein 1 (AP-1) family members cooperate with hepatocyte-specific transcription factors, including HNF4A, in maintaining cellular functionality of mature human hepatocytes. Comparative analysis identified distinct transcription factor sets specifically active in human hepatocytes and organoids. Amongst these ELF3 is unique to intrahepatic cholangiocyte organoids and its expression level negatively correlate with expression of hepatic marker genes. Functional analysis of ELF3 furthermore revealed that *ELF3* depletion optimizes the formation of hepatocyte-like cells from intrahepatic cholangiocyte organoids.

**Conclusions:** Collectively, our integrative analysis provides insights into the transcriptional regulatory networks of human hepatocytes and hepatic organoids, thereby informing future strategies for better establishment of urgently-needed hepatic model systems *in vitro*.

## Introduction

The human adult liver is a complex and regenerative organ that is composed by parenchymal and non-parenchymal cell types. Primary human hepatocytes (PHHs), which make up more than 80% of the total liver mass, play a major role in supporting a wide range of liver functions including glycogen storage, lipid/carbohydrate metabolism, and exogenous/endogenous compounds inactivation (1). Fully matured PHHs are functionally heterogeneous along the porto-central axis of the liver lobule, known as liver zonation. PHHs stay in a quiescent state but can re-enter the cell cycle to start proliferation following damage (2,3). Hepatocyte regeneration is mainly separated into three phases: the priming phase, the proliferation phase, and the termination phase (4). Studies on liver regeneration has mainly been performed using mouse models to investigate the original location of the initiation of hepatocyte regeneration. However, this process remains unclear in humans due to the lack of a proper model.

Long-term expansion of functional human hepatocytes *in vitro* is currently the main model for studying various aspects of hepatocytes, such as drug toxicity prediction, modeling liver diseases for clinical studies, and transplantation. However, the utilization of PHHs is limited due to the difficulty of maintaining their functional and cellular identity and lack of their capacity to proliferation *in vitro*. Therefore, numerous studies in the last decades have aimed to solve these bottlenecks, and currently one of the most promising approaches is the use of 3D *in vitro* cultures, liver organoids (5). Different from cells in 2D culture systems, organoids form a 3D structure by self-organization, display high levels of genetic stability and recapitulate certain structural and functional aspects of the organs *in vivo* (6,7). Human liver organoids can be generated from various sources including induced pluripotent stem cells (iPSCs), or from fetal and adult primary tissues (5). Depending on their origin, organoids display distinct characteristics, functions and differentiating/self-renewing capacities (5). For example, human adult hepatocyte organoids were recently generated that display both progenitor and mature hepatic features. However, the establishment and maintenance of these hepatocyte-derived organoids remain challenging (8,9). Human intrahepatic cholangiocyte organoids (ICOs), generated from EPCAM positive bile duct cells and maintained in expansion medium (EM; i.e. WNT pathway activated), are on the other hand, highly proliferative and can be differentiated toward hepatocyte-like cells when cultured in the hepatocyte differentiation medium (DM) (6,7), making it as an attractive model for long-term maintenance of liver cells *in vitro*. However, hepatocyte-like cells differentiated from ICOs remain immature compared to PHHs, which hampered the usage of this model in therapeutics and fundamental studies (10). Thus, there is an urgent need for further improvement of functional ICO-derived hepatocyte differentiation *in vitro,* for which further understanding of the gene regulatory mechanisms underlying hepatocyte differentiation and maturation is highly beneficial.

Gene expression patterns determine cell identity, and they are accurately controlled by gene regulatory networks (GRNs) of transcription factors (TFs), epigenetic modifiers, and chromatin remodelers. In the liver, studies have shown crucial roles of hepatic TFs in controlling multiple biological processes, and dysregulation of hepatic TFs has been associated with developmental defect, failure of liver regeneration and various liver diseases (11–13). Therefore, a better understanding of how TFs orchestrate gene expression in hepatocytes would provide important insights for the molecular mechanism of hepatocyte maturation both *in vivo* and *in vitro*. TFs required for cell fate decision during liver development have been explored mostly in mouse models or during *in vitro* pluripotent stem cell differentiation. For example, TFs in the hepatocyte nuclear factor (HNF) family, such as HNF4A, FOXA1 and FOXA2 (also known as HNF3A and HNF3B), together with GATA-binding factor (GATA) 4/6 and CCAAT enhancer binding protein (C/EBP) A/B are liver-enriched TFs that are important for normal liver development (14). However, substantial species variability between mouse and human limits the use of the mouse models for understanding human liver biology and pathogenesis. Moreover, the gap between human liver organoids and mature hepatocytes is still to be explored. It remains unclear whether other TFs are involved in the regulatory network, especially in PHHs, and how the gene regulatory mechanisms in PHHs compare to that in ICO-derived hepatic like cells.

In this study, we combined RNA-seq at single cell resolution and ATAC-seq to dissect the gene regulatory mechanisms in PHHs and ICOs. Using an optimized two-step perfusion protocol, we isolated a nearly pure population of PHHs. scRNA-seq analysis displayed that the heterogeneity in PHHs obtained is driven by the liver zonation. Integrative analysis in PHHs demonstrate that Activator protein 1 (AP-1), dimeric transcription factors composed of Jun, Fos or ATF (activating transcription factor) known as key TFs in the hepatic regeneration initiation (24), contribute to the establishment of the hepatic gene expression program in conjunction with typical liver specific transcription factors, for example HNF4A. Comparison of gene regulatory analysis revealed distinct TF sets required for PHHs and ICOs, while surprisingly gene regulatory TFs and chromatin accessibility in EM-ICOs and DM-ICOs only showed subtle differences. Further study on ELF3, the most prominent TF that is specifically expressed in ICOs, showed that depletion of *ELF3* during hepatocyte differentiation of ICOs promotes expression of mature hepatocyte markers, suggesting that ELF3 functions as a barrier of hepatocyte differentiation in ICOs.

## Materials and Methods

### Ethics statement

Primary human liver cells were freshly isolated from patients undergoing liver surgery. General approval for use of remnant, anonymized surgical material was granted in accordance to the Dutch ethical legislation as described in the Medical Research (Human Subjects) Act and confirmed by the Committee on Research involving Human Subjects, in the region of Arnhem-Nijmegen, the Netherlands (CMO-2019-5908). Metadata associated with these samples can be found in Table S3. The use of liver biopsies for research was approved by the MEC of the Erasmus MC (MEC-2014-060) and by the CMO-Radboudumc (CMO-2015-2062).

### Human liver tissue dissociation and single cell isolation

Human liver specimens obtained from liver resections were used for primary hepatocyte isolation following the two-step perfusion protocol as described before (32). Briefly, the liver segment was first perfused with HBSS (Thermofisher, 14170-138) containing 0.64 mM EDTA (Thermofisher, 15575-038) and 10mM HEPES (Thermofisher, 15630-056) to wash away the blood, and then perfused with HBSS supplemented with 10mM HEPES to remove any residual EDTA. Next, the liver segment was perfused with HBSS supplemented with 10mM HEPES, 0.75 mg/ml CaCl2 and low concentration of collagenase VIII (3333 units per 50ml), followed by the perfusion using the same buffer but with high concentration of collagenase VIII (13,333 units per 50ml) for about 30 min until the liver tissue became soft. All the perfusion steps were performed at 37 ⁰C. After the collagenase dissociation, DMEM (Thermofisher 31885-023) with 10% FBS was used to inactivate the collagenase activity, and liver tissue was cut into small piece to release cells into the DMEM medium. The resulting cell suspension was collected and filtered with a 100µm cell strainer. The cells were pelleted at 50g for 5 min. To remove non-hepatocytes, cells were washed with DMEM and pelleted 5min at 50g for two more times (until the supernatant is clear after the centrifugation). To remove the nonviable hepatocytes, the cell pellet was resuspended in 28% percoll and centrifuged at 150g for 20 min. Cells were counted with trypan blue, and percoll gradient centrifugation were repeated until the percentage of dead cells is about 20%. The isolated hepatocytes were directly used for single cell sorting.

### Intrahepatic cholangiocyte organoid (ICO) culture, differentiation and single cell isolation

ICOs (n=3) were obtained from Erasmus MC and from the biobank at department of Gastroenterology and Hepatology, Radboudumc (Nijmegen, NL). To initiate ICOs, tissue biopsies from liver (circa 0.5-3 cm2, n=1) were obtained from donor livers during liver transplant procedures performed at the Erasmus MC, Rotterdam, the Netherlands. ICOs were cultured and differentiated towards hepatocyte-like cells as published before (6,27). Briefly, ICOs were cultured in Basement Membrane Extract (BME, Trevigen, 3533-010-02) and the expansion medium was refreshed every 3 days. For the hepatocyte differentiation, the organoids were first cultured in expansion medium supplemented with BMP7 (PEPROTECH, 120-03P) for 5 days, and then switched to differentiation medium for more than 10 days. To obtain single cells, BME was removed by 10-fold dilution in cold DMEM/F-12 (Thermo, 12634-010). The organoids were then digested using 0.05% Trypsin-EDTA (Thermo, 25300-054) until 80-90% of the cells were single cells. Trypsin was removed and cells were collected for further use.

### Immunofluorescence for organoids

BME was first removed using Cultrex Organoid Harvesting Solution (Trevigen, 3700-100-01), and ICOs were collected. Cells were fixed with PFA 4% on ice for 30 min and then washed 3 times with cold PBS. After fixation, cells were blocking using blocking buffer (0.5% Triton-1000, 1% BSA, 1% DMSO in PBS) at room temperature for 3 hours. Then primary antibodies (EPCAM (eBioscience, 14-9326-82) used in 1:100 and ALB (Bethyl Laboratories, A80-229A) used in 1:50) were diluted and incubated with cells at 4 ⁰C for more than 24h. After 3 times washing with PBS + BSA 1%, cells were incubated with secondary antibodies (Donkey anti-Goat 594 (Thermo Fisher Scientific, A32758) and Donkey anti-Goat 488 (Thermo, A-21202) both used in 1:250) for 2 hours at room temperature. Cells were washed 3 times with PBS + BSA 1% and stained with Hoechst 33342 (Thermo, 62249) for 1 hour at room temperature. Finally, cells were mounted in the chamber with the mounting buffer (IBIDI, 50001) and imaged by Leica SP8 confocal microscope. Images were further processed using ImageJ (66).

### siRNA transfection

At day 5 of ICO differentiation, cells were collected. For transfection, siRNAs (negative control No.1 siRNA, Thermo, 4390843; siELF3-s4623 and siELF3-s4624, Thermo, 4427037) with a final concentration of 40 nM and Lipofectamine 3000 (Thermo, L3000015) reagent were used following the manufacturer’s protocol (27). To obtain efficient knock down, two separate siRNAs targeting *ELF3* were combined. After transfection, cells were seeded in BME and cultured in differentiation medium supplied with 10 µM Rock inhibitor (Gentaur, A3008) for the first 3 days. Transfected cells were then harvested for RNA isolation at day 8 and day 12 after differentiation.

### RNA extraction and RT-qPCR

Cells collected from either 3D culture organoids or sorted cells were lysed in RLT buffer from RNeasy kit (Qiagen, 74106). RNA samples were isolated using the same kit following the manufacturer’s recommendations. 100-300 ng of RNA was used for cDNA synthesis using iScript cDNA Synthesis Kit (Bio-Rad, 1708891) following the manufacturer’s manual. RT-qPCR was performed in triplicate on a CFX96 Real-Time System (Bio-Rad) with SYBR Green Supermix (BIO-RAD, 1725006CUST). Data was normalized to *GAPDH*. Primer sequences are provided in Table S4.

### ATAC-seq library preparation

25000 cells were washed once with cold PBS, and then resuspended in Tn5 reaction mix (12.5 µl 2x TD buffer (homemade), 1.25µl Tn5 enzyme (homemade), 0.5 µl Digitonin (Promega, G9441), and 11 µl H_2_O). Cells were incubated in the thermo mixer at 37 ⁰C, 800 rpm for 20 min. After the incubation, reaction cleanup was performed using QIAGEN MinElute Reaction Cleanup Kit (28204). The purified tagmented DNA samples were barcoded with Nextera Index Kit primers and amplified via PCR followed by a AMpureXP bead (Agencourt, A63882) size selection. Briefly, after first 5 PCR cycles, 5ul of PCR reaction was used for qPCR to determine the number of additional cycles needed. After amplification, PCR reaction was cleaned up with 0.65x AMpureXP beads followed by 1.8x SPRI beads to select DNA fragments from 150 to 600 bp. Finally, libraries were quantified on the bio-analyzer using DNA high-sensitivity kit, and sequenced on an Illumina NextSeq 500 with 38bp paired-end reads at a depth of about 20 million reads per sample.

### Single cell cDNA library preparation

Single-cell RNA libraries were constructed according to the mCEL-seq2 protocol (67). Briefly, 100 nL lysis buffer (0.2% Triton X-100, 2mM dNTPs, 1:50.000 ERCC ExFold RNA spike-in (Thermo, 4456739)) was added per well using the Nanodrop Ns-2 Stage (BioNex). For the *in vitro* reverse transcription reaction, SuperScript™ II Reverse Transcriptase (Invitrogen, 18064-14) was used with the following the protocol, 4 ⁰C for 5 min; 25 ⁰C for 10 min; 42 ⁰C for 60 min; 70 ⁰C for 10 min. Next, 960 nL second-strand reaction mix containing *E. coli* DNA ligase (NEB, M0205L), *E. coli* DNA polymerase I (NEB, M0209L) and Random hexamer (sigma, 11034731001) were added per well for second-strand synthesis following the protocol, 16 ⁰C for 2 hrs. All the cDNA from one 384-well plate was pooled. *In vitro* transcription was performed using the MEGAscript™ T7 Transcription Kit (Invitrogen, AMB 1334-5) by incubating the reaction at 37 ⁰C for 14 hrs. The amplified RNA was fragmented and used for cDNA synthesis using Superscript II and the random octamer primer. Libraries were amplified for 9 cycles for the hepatocyte libraries and 8 cycles for the liver organoid libraries using Phusion® High-Fidelity PCR Master Mix (NEB, M0531S) with the Nextflex primers, and quantified on the bio-analyzer using DNA high-sensitivity kit (Agilent, 5067-4626). Barcoded libraries were sequenced on an Illumina NextSeq 500 with 38bp paired-end reads at a depth of about 25 million reads per plates.

### scRNA-seq data analysis

Barcode information was extracted from read 1 and added to the ID of paired read 2 using UMI-tools 1.1.0 (68). The barcoded reads containing unique molecular identifier (UMIs, 1-8bp) and cell barcode (9-16bp) were not used for mapping. Only the paired read 2 were aligned to a modified human genome (hg38) containing 92 ERCC spike-in sequences using STAR-2.7.6a with default settings (69). After mapping, featureCount 2.0.1 was used to assign the reads to genes (70). The number of UMIs per gene was counted for each cell and the gene-cell count tables were generated using UMI-tools 1.1.0 with the following parameters, --per-gene, --per-cell. For the downstream analysis, Seurat 3.2.3 was applied for data filtering, batch correction, normalization, clustering, visualization and differentially expressed gene analysis (71). Briefly, Seurat objects were created per donor using gene-cell matrices. Genes detected (UMIs > 0) in at least 5 cells were kept. Cells that are with less than 1000 UMIs or more than 50% of mitochondrial gene expression (mt%) were removed. UMI counts were then log2 transformed and normalized to sequencing depth per cell, and number of total UMIs and mt% were regressed out during the scaling to correct for sequencing depth and mt%.

For scRNA-seq data analysis within PHHs or ICOs, to minimize the batch effect, Seurat objects from different donors were integrated using the functions FindIntegrationAnchors and IntegrateData. Next, Top 2000 variable genes were found and used for PCA analysis with the setting, npcs=50. Seurat function FindNeighbors, FindClusters, and RunUMAP were used to construct a Shared Nearest Neighbor (SNN) Graph, identify clusters of cells based on the SNN graph, and finally to reduce the dimension for the visualization. Finally, differentially expressed genes from each cluster were identified using Wilcoxon Rank Sum test, and the significant ones (adj-p<0.05) were used for pathway analysis via Enrichr web tool (72,73).

For integration analysis of PHHs and ICOs, PHH and ICO Seurat objects were merged using Seurat merge function. To compare ICOs to PHHs only, non-hepatocyte cluster in the PHHs dataset was not included in the integration. Then the newly merged Seurat object was first scaled with regressing out the number of total UMIs, mt% and biological replicates, and then used for further analysis following the same procedure as mentioned before.

### Correlation analysis with zonally-expressed human and mouse transcripts

For correlation analysis of zonally-expressed genes between human and mouse hepatocyte, we used a set of 94 genes previously identified to be expressed in both human and mouse (16,17). Average expression levels of each gene were calculated for all the 4 hepatocyte clusters (from this study) and then scaled and centered by z-scores. Same analysis was applied for nine layers of mouse hepatocytes (17). Pearson correlation was calculated to compare the 4 human hepatocyte clusters with the nine layers of mouse hepatocytes.

### Cell type prediction analysis of ICOs

To predict the cell type of the cells in ICOs, we used two published scRNA-seq datasets generated from the human liver as references (16,28), extracted the data of hepatocytes from all zones and cholangiocytes, and trained a cell type’s Random Forest classifier by Python package sklearn (v.0.24.2). Only the top 5000 variable genes were using for the training. Next, ICOs from both culture conditions were classified using this classifier, and assigned to a cell type in the reference for each single cells. Finally, sankey plots were generated based on the assignments using R package networkD3 (v.0.4).

### Transcription factor regulon analysis

SCENIC R version (74) was used for gene regulatory network analysis. Briefly, raw gene-cell count matrix was extracted from “RNA” assay in Seurat object and used as input for SCENIC. The matrix was log 2 transformed, and co-expression network was calculated using runGenie3 with default setting. To predict the transcription factor regulons, *hg19-tss-centered-10kb-7species.mc9nr.feather* motif database from RcisTarget was used. Regulons were created with the following parameters, minGenes = 20, coexMethod=“top5perTarget”. The Area Under the recovery Curve (AUC) scores were generated per cells and imported into Seurat for downstream analysis and visualization.

### ATAC-seq and ChIP-seq analysis

For ATAC-seq data analysis, pair-end reads were mapped to the human reference genome (hg38) using BWA 0.7.17-r1188 mem (77) with default settings. Next, duplicates and reads with mapping quality lower than 30 were removed using samtools 1.7 (75). Open chromatin profile tracks were generated using deeptools 3.5.0 (78) bamCoverage function with the following setting, -- normalizeUsing CPM. For peak calling, MACS2 v2.2.7.1 (79) was used with the following settings, -g hs, --nomodel. Overlapping and unique peak sets between biological replicates or different samples were identified using homer v4.11.1 (80) mergePeaks function with the parameter, -d given. To minimize the donor variation and get most confident peak sets, only the peaks presented from both biological replicates were used for further analysis. For spearman correlation analysis, ATAC peaks were first merged from all the samples, and the ATAC signal was quantified in all peaks from each sample using bedtools v2.29.2 multicov function (81). The peak-sample count table was either imported into DEseq2 1.26.0 (76) for making the PCA plot following the online tutorial, or CPM normalized for spearman correlation analysis using rcorr function from R package Hmisc 4.4-2. Genes targeted by HNF4A, JUND or both proteins were selected according the following criteria, expression level higher than 100 UMIs from all summed single cells and the binding sites detected within TSS±5kb of the genes. Next, HNF4A and JUND targeted genes were used for GO analysis using Enrichr web tool. For motif analysis, GimmeMotifs v0.15.1 (82) was applied using 200bp around peak summit regions as input with the following settings, -s 0, -b gc. For differentially enriched motif analysis, the unique peak sets from each sample were merged. Then normalized read coverage was quantified (as CPM) in each peak region (200bp around summit point) for each sample. After log2 transformation, the matrixes were imported into GimmeMotifs (82), and maelstrom function was used with the default settings.

For ChIP-seq analysis, the ChIP-seq profiles were downloaded from ENCODE (83–85). Read mapping, filtering and peak track generation were done using the same strategy as ATAC-seq analysis. MACS2 was used for peak calling with the following settings, -g hs. For spearman correlation analysis, ChIP-seq signal was quantified for each TFs in the ATAC peak regions in PHHs. After CPM normalization, the count table was imported into R and correlation was calculated using the same strategy as ATAC-seq spearman correlation analysis. The intersecting peak regions were identified using homer mergePeaks with the parameter, -d given. For the motif analysis, GimmeMotifs was applied with the settings as mentioned before (82).

### Quantification and statistical analysis

For studies where statistical analyses were performed, at least three biological replicates were used unless otherwise indicated. For two-condition comparisons, two tailed student t-tests were used with a p-value <0.05 as the threshold for significance. For multiple testing, a Wilcoxon Rank Sum test was used in Seurat (as default) and a Wald significance test was used in DEseq2 (as default) for differentially expressed gene determination with an adj-p<0.05 as the threshold for significance. For pathway and GSEA analysis, Fisher exact test and Benjamini-Hochberg method were used as suggested with an adj-p<0.05 as the threshold for significance.

## Results

### Liver zonation is the main driver of the heterogeneity among primary human hepatocytes in situ

Previous scRNA-seq datasets were mainly generated from the whole liver, which is aiming to recapitulate the full complexity of cell types within the liver tissue (15,16). However, unbiased isolation of all cell types leads to a limited and inconsistent proportion of primary human hepatocytes (PHHs) from each donor/sample hence compromising in-depth analysis of PHHs. Therefore, we adapted a two-step perfusion isolation approach to obtain pure human hepatocytes for single-cell RNA-seq analysis (see Materials & Methods and Figure 1A). We obtained 2528 high-quality cells from three adult livers, and identified five different clusters after normalization and batch correction (Figure 1B-C and Figure S1A-B). Conforming the efficiency of our PHH isolation protocol, the majority of the cells displayed high expression levels of matured hepatocyte markers but low or no expression of liver progenitor cell- or cholangiocyte-associated genes (Figure S1C). In fact, only about 2% of the cells (cluster 5) represented other cell types and showed expression of well-known markers for endothelial/epithelial and immune cells (*DCN*, *CD74*, *ZEB2* and *TCF4;* Figures S1C-S1D, and Table S1), which we excluded from further analysis.

**Figure 1.**
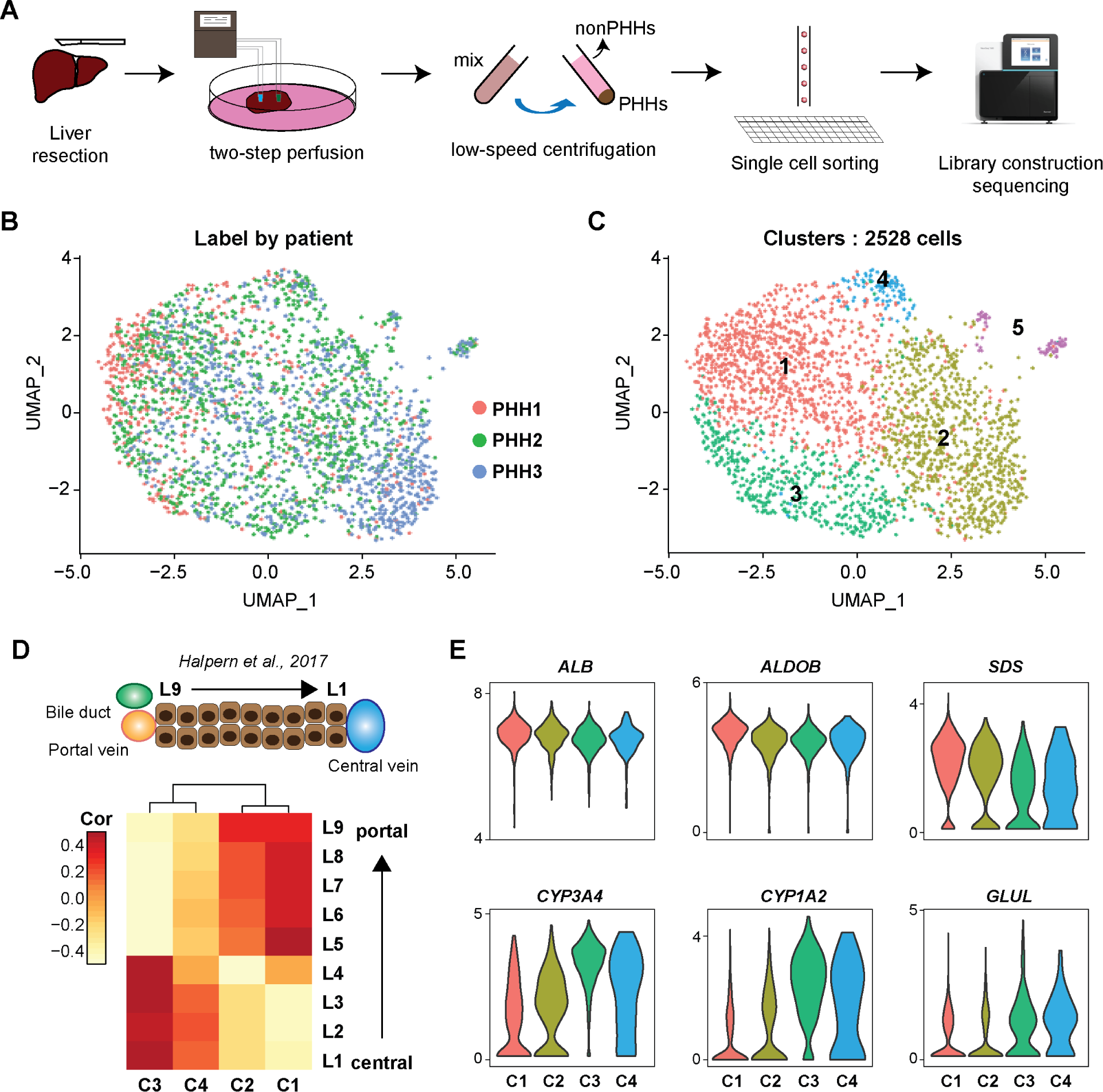
scRNA-seq analysis shows that liver zonation is the main driver of heterogeneity among human PHHs. **(A)** overview of single-cell isolation from liver segments following a modified two-step perfusion protocol and FACS sorting workflow. **(B-C)** UMAP plots of scRNA-seq data colored by donors (left) and cluster number (right). **(D)** Heatmap showing the correlation of selected zonal genes between hepatocyte clusters identified in this study and the nine layers of mouse liver lobule (L1, central; L9 periportal). **(E)** Violin plots showing expression of representative zonal markers (periportal: *ALB*, *ALDOB*, and *SDS*; pericentral: *CYP3A4*, *CYP1A2*, and *GLUL*) across hepatocyte clusters.

Genes most prominently enriched in cluster 1 include *SDS* and *ASS1,* while *CYP1A2*, *CYP3A4*, *GLUL* are enriched in cluster 3 (Figure S1D), all representing markers known to be associated with liver zonation. Indeed, when comparing with zonally resolved expression data from mouse, we observed a clear zonal pattern with clusters 1 and 2 behaving more similar to periportal zone and clusters 3 and 4 sharing pericentral features (16,17) (Figure 1D-1E and Figure S2). Altogether, our data further strengthen that porto-central zonation is the primary driver of heterogeneity amongst PHHs.

### Gene regulatory network analysis identifies crosstalk between AP-1 and liver-specific transcription factors in mature hepatocytes and during hepatocyte regeneration

Transcription regulation determines cell identity and function. To identify the key regulatory factors in the primary hepatocytes, we applied single-cell regulatory network inference and clustering (SCENIC). SCENIC identifies potential master TFs as well as their downstream target genes (called regulons) by integrating both co-expression modules between TFs and target genes and *cis*-regulatory motif enrichment at these target gene loci. Using hepatocytes from cluster 1 to 4, we detected in total 63 regulons with relatively little difference between the clusters (Figure S3A). We identified HNF4A, CEBPB, CEBPD, and ONECUT2 regulons, of which the corresponding transcription factors are well-known and required during liver development (14,21) (Figure 2A). FOXO1 and FOXP1 TFs regulates hepatic glucose homeostasis through response to insulin (22,23). While we observed subtle difference of regulons between different clusters, in general we found that the regulons of TFs from AP-1 family such as JUND, JUN, JUNB, FOS, FOSB, EGR1 and ATF3, classically associated with cellular proliferation and oncogenesis, are highly active in all primary hepatocytes, which is even more pronounced in cluster 4 (Figure 2A). Taking a closer look, we found that genes enriched in cluster 4 include not only genes in AP-1 family, but also *TNFAIP3*, *GADD45B*, *SERPINE1*, and *G0S2* (Figure S3B) known to be up-regulated immediately in the priming phase of mouse hepatocytes as an initial step of the hepatic regeneration (24,25). Thus, cells in cluster 4 might represent the elusive human hepatocytes in their priming phase. Overall, our analysis indicates that besides liver regeneration, AP-1 TFs could also play an important regulatory role in all mature hepatocytes.

**Figure 2.**
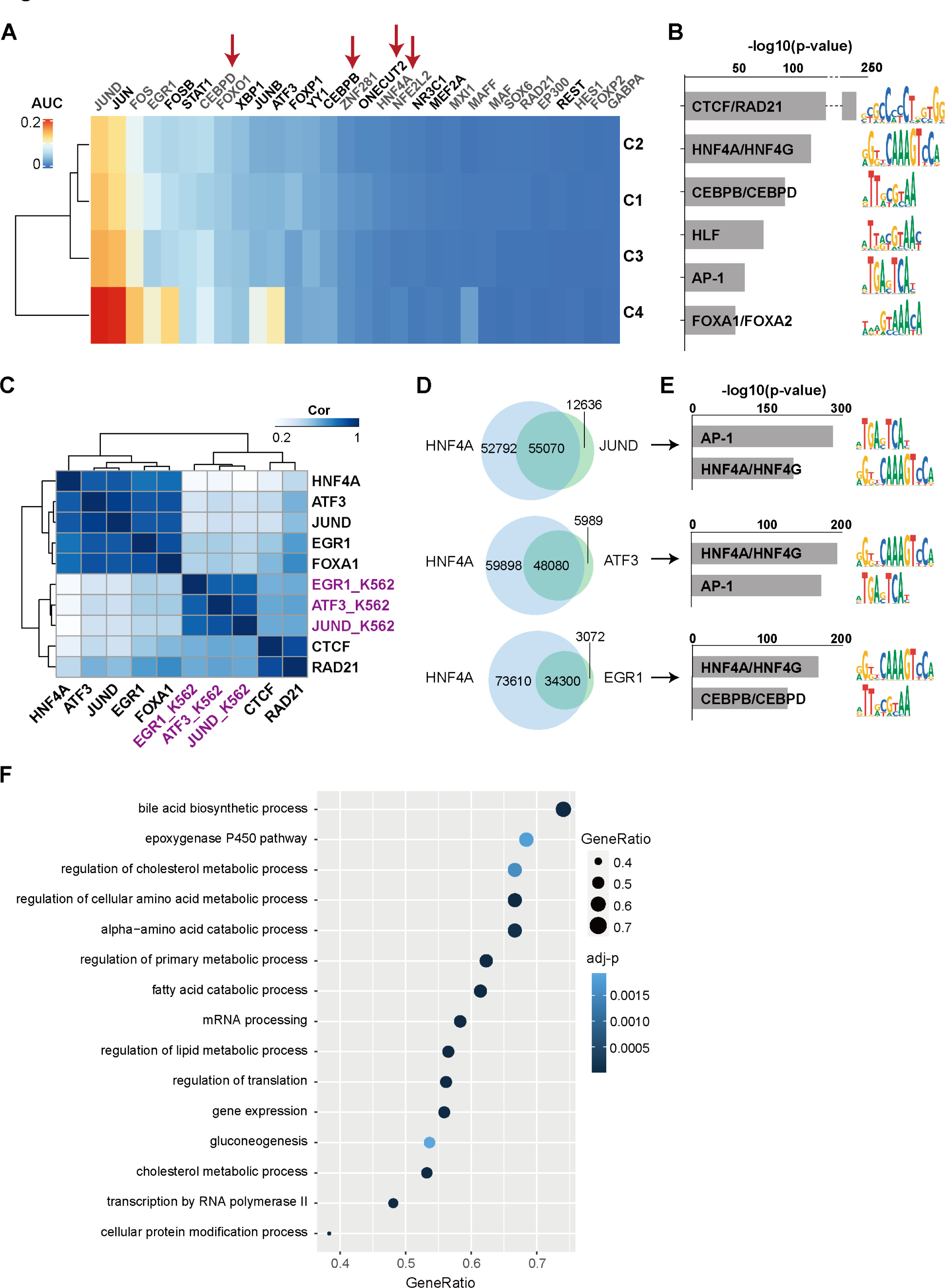
Gene regulatory network analysis identifies crosstalk between AP-1 and liver-specific transcription factors in PHHs. **(A)** Heatmap of enrichment of regulons (shown as average AUC score) identified in the 4 hepatocyte clusters. Regulons in grey represent regulons in which non-direct motifs were used for prediction of potential targets of the TFs, and the black ones represent regulons in which direct motifs were used for prediction of potential targets of the TFs (see Materials & Methods). Red arrows point at the well-known hepatic TFs. **(B)** List of Top 6 motifs enriched in ATAC peaks in PHHs. Motifs are ranked by −log10(p-value). **(C)** Heatmap of spearman correlation matrix of normalized TF binding intensities in PHHs open chromatin regions identified by ATAC-seq. TFs labeled in black are from liver and ones in purple are from K562 lymphoblast cell line (as control). **(D)** Venn diagram showing the intersection between binding sites of HNF4A and those of JUND, ATF3 and EGR1, respectively. **(E)** The top 2 motifs enriched in overlapping ChIP-seq peaks regions of HNF4A and JUND, ATF3 and EGR1 (as in D), respectively. Motifs were ranked based on −log10(p-value). (**F**) Dot plot of GO analysis (biological process) showing the selected significant (adj-p<0.05) pathways enriched for the HNF4A and JUND co-bound genes. Circle color indicates the significance of the enrichment and the size represents the gene ratio.

In parallel to our scRNA-seq analysis we carried out ATAC-seq on the PHHs isolated from 2 different donors and performed motif analysis on the open chromatin regions. Here, we detect enrichment of CTCF/RAD21, HNF4, CEBP, AP-1 and FOXA1 motifs in the accessible chromatin regions (Figure 2B), in line with the SCENIC scRNA-seq analysis. To further investigate the roles of AP-1 TFs in hepatocytes, we reanalyzed published JUND, ATF3, EGR1 FOXA1, HNF4A, CTCF and RAD21 ChIP-seq datasets from human liver tissue and integrated them with our ATAC-seq data (from ENCODE), including JUND, ATF3 and EGR1 ChIP-seq datasets from K562 lymphoblast cell line as controls (from ENCODE). In agreement with the motif analysis, high enrichment of HNF4A and AP-1 (EGR1, JUND, and ATF3) TF occupancies were found in the ATAC peaks in PHHs (Figure S3C). We evaluated and compared genome-wide binding profiles of the selected TFs in accessible regions in the PHHs by quantifying ChIP-seq reads within PHH ATAC-seq peak regions. Spearman correlation analysis of TF binding within open chromatin regions revealed that AP-1 TF binding profiles displayed strong correlation with HNF4A and FOXA1 binding profiles in the liver, but not in K562 lymphoblast cells (Figures 2C and S3D). We identified ChIP peaks for HNF4A, JUND, ATF3, and EGR1 to determine the overlap in occupancies. This analysis showed that the 3 AP-1 TF binding sites are highly enriched in HNF4A peak regions (Figure S3C). 81% (55070 of 67706), 89% (48080 of 54069), and 92% (34300 of 37372) of JUND, ATF3 and EGR1 peaks are intersecting with HNF4A binding sites, respectively (Figure 2D). Accordingly, motif analysis revealed a noticeable enrichment for HNF4A motif at JUND, ATF3, and EGR1 peaks (Figure 2E). To elucidate the potential regulatory roles of AP-1 TFs in PHHs, we next focused on genes that are targeted by HNF4A and JUND. Among 7716 genes with detectable levels of transcript abundance in our scRNA-seq data, we observed many genes (5137) that are bound by both HNF4A and JUND compared to target genes bound by either HNF4A (743 genes) or JUND (62 genes) only (Figure S3E). These HNF4A/JUND co-occupied genes showed significant enrichment of Gene Ontology categories associated with fundamental biological pathways, including gene expression, RNA processing, protein modification, transcription regulation as well as multiple metabolic processes such as lipid, fatty acid, bile acid metabolic processes, epoxygenase P450 pathway, and gluconeogenesis (Figure 2F). Altogether, our analysis strongly indicates that AP-1 TFs, by cooperating with liver specific TFs, are associated with both fundamental cellular and liver-specific functions in hepatocytes. Furthermore, AP-1 TFs, as shown in mouse (24), are also prominent features of regenerating human hepatocyte in their priming phase.

### Integrative analysis uncovers distinct TFs required in PHHs and ICOs, and identify ELF3 as a prominent TF in ICOs

Previous studies demonstrated that ICOs have the potential to differentiate towards the hepatocyte lineage by culturing them in differentiation medium (DM) (6,26). However, these organoid-derived hepatocyte-like cells are not fully matured as compared to primary hepatocytes. To identify the gene regulatory differences among ICOs in different culture conditions as well as between ICOs and PHHs that could explain the above observations, we carried out single cell RNA-seq on the ICOs cultured in either EM or DM (EM-ICOs vs DM-ICOs). Differentiation of liver organoids was performed with 3 organoid lines derived from different donors as previously described (27) (Figure S4A). In total, we generated single-cell transcriptomic profiles with high quality (UMIs>1000, and mt%<50%) from 1268 ICO cells. UMAP indicated cells are clustered based on culture conditions (Figure S4B). Cell cycle analysis showed more DM cells in G1 phase compare to EM cells, suggesting that DM-ICOs are, as expected, less proliferative (Figure S4C). In line with previous reports (6,26), we detected a down-regulation of WNT target gene *LGR5* and progenitor/cholangiocyte marker *SOX9*, and up-regulation of hepatocyte markers (*ALB*, *CYP3A4*, *GLUL* and *GC*) (Figures S4D-S4F). During differentiation, the DM-ICOs do not show a clear gene expression signature associated with either pericentral or periportal zone (Figure S4G). Expression of cholangiocyte markers (*TFF1*, *TFF3*, and *MUC5B*) in DM-ICOs decreased compared to EM-ICOs, while other cholangiocyte markers such as *EPCAM*, *KRT7* and *KRT19* remained largely similar (Figure S4F), indicating that ICOs under hepatocyte differentiation condition still retain cholangiocyte features. Next, to better understand the transcriptome of both EM- and DM-ICOs, we built a Random Forest classifier based on the published human liver scRNA-seq data (16,28), and evaluated the similarities between ICOs and the *in vivo* human liver cells (see Materials & Methods). We found that ICOs, independent of culture conditions, substantially resembled human cholangiocytes (Figure S4H). Therefore, our analyses indicate that ICOs under both expansion and hepatocyte differentiation conditions largely retain cholangiocyte features.

To gain insights into the gene regulatory mechanisms that could explain the lack of full maturation of ICOs, we performed integrated analysis of the ICO and PHH datasets. We found that cells were clearly separated based on the different cell types with clear distinct transcriptional signatures of PHHs and ICOs (Figure 3A). Next, we used SCENIC to identify master TFs in hepatocytes and ICOs from both conditions. In total, 56 regulons were found, some of which were enriched across different cell types such as AP-1 TFs (JUN, JUND, FOS, FOSB, and ATF3), STAT1, YY1, XBP1, and NFE2L2. Some TFs were specific to either PHHs or ICOs (Figure 3B). For example, MLX Interacting Protein Like (MLXIPL), also known as ChREBP, is a glucose-sensitive transcription factor that is crucial for glucose metabolisms and *de novo* lipogenesis (29). Peroxisome proliferator-activated receptor gamma coactivator 1-alpha (PPARGC1A) is required for nutrient metabolism in the liver (30,31). Both TFs are associated with metabolism and are more enriched in PHHs, in line with the fact that PHHs display a more physiologically relevant gene expression pattern as compared to ICOs. On the other hand, we identified ELF3 and EHF as main regulons in ICOs (EM and DM) that are absent in PHHs. In line with the enrichment of regulons, expression levels of these TFs also displayed similar trends between PHHs and ICOs (Figure 3C).

**Figure 3.**
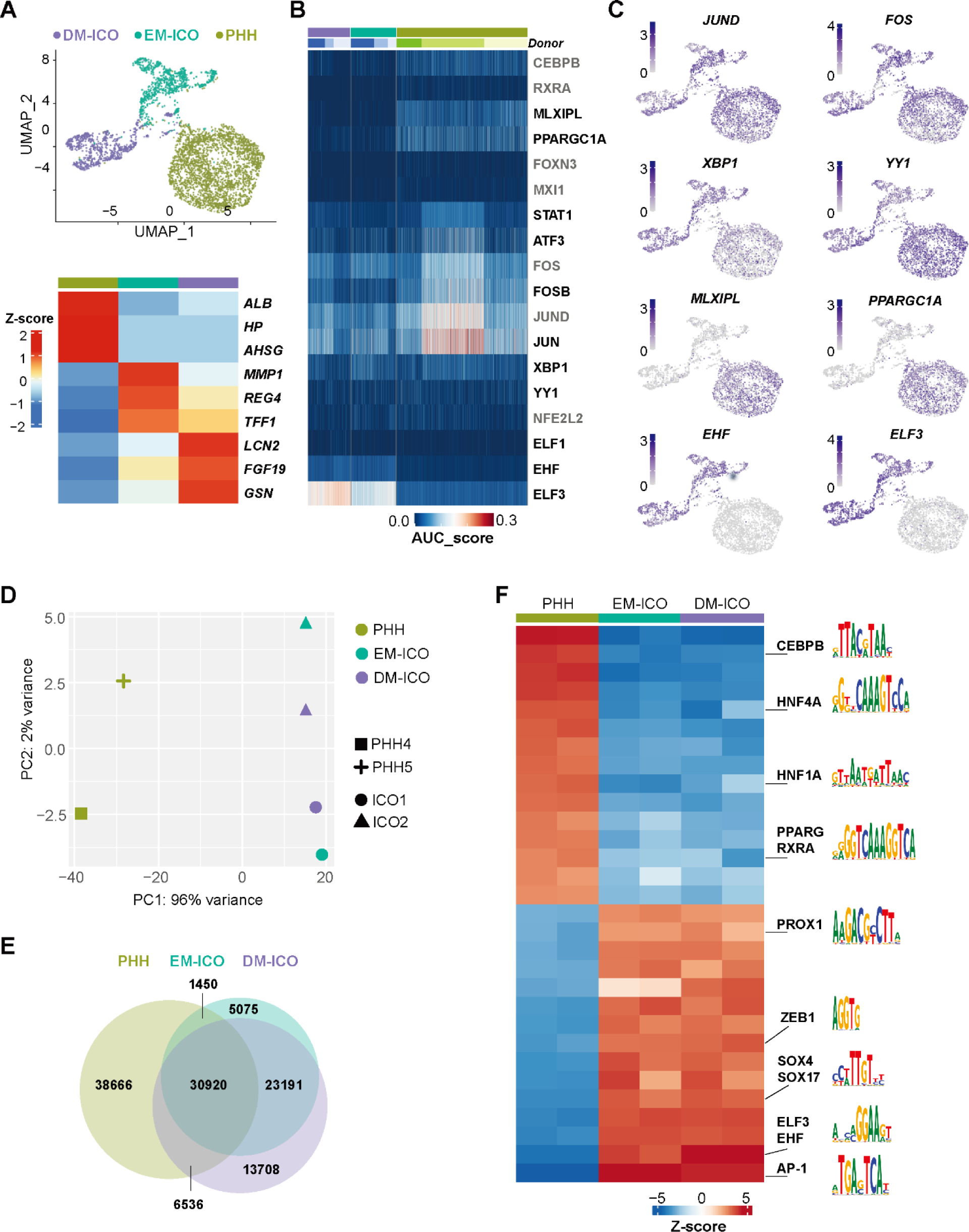
Integrative analysis of scRNA-seq and ATAC-seq identifies distinct TFs required in PHHs and ICOs. **(A)** UMAP of scRNA-seq data from PHHs and ICOs maintained in EM and DM (up panel) and heatmap showing the expression of top 3 marker genes in Z-score format for each cell type (bottom panel). Cells are colored based on the cell types. **(B)** Heatmap of the AUC score of selected regulons in each single cell from PHHs and ICOs as analyzed by SCENIC. Colors of each cell types match those of (A). **(C)** Expression UMAPs of *JUND*, *FOS*, *XBP1*, *YY1*, *MLXIPL*, *PPARGC1A*, *EHF*, and *ELF3*. Data was normalized to sequencing depth and shown in log2 format. **(D)** PCA plot showing the similarities of the genome-wide chromatin state between PHHs, EM-ICOs, and DM-ICOs. Biological replicates are shown in different shapes, and cell types are shown in different colors as in (A). **(E)** Venn diagram showing the intersection between ATAC peaks of PHHs, EM-ICOs and DM-ICOs. **(F)** Heatmap showing motif enrichment analysis between PHHs, EM-ICOs, and DM-ICOs in the Z-score format using Gimme Motif maelstrom. Motifs of interest are shown on the right.

To explore the chromatin landscape in ICOs, we performed bulk ATAC-seq on two ICO lines cultured in EM and DM, and examined the divergence of chromatin accessibility profiles between PHHs, EM-ICOs and DM-ICOs. Overall, we observed little difference between biological replicates and culture condition of ICOs (EM and DM) but substantial difference between PHHs and organoids (Figures 3D and S5A-C). When comparing ATAC peaks across different samples, more shared ATAC peaks were found among ICOs than those between PHHs and ICOs (Figure 3E), in agreement with the correlation analysis. ATAC signal over top 2000 unique peaks from each sample also show clear difference between PHHs and ICOs (Figure S5B). Therefore, our analysis highlights distinct chromatin landscape between PHHs and ICOs.

TFs bind to DNA and regulate gene expression through specific DNA motifs. Thus, we performed motif analysis using GimmeMotifs to determine which TFs were potentially associated with open chromatin regions from EM-ICOs and DM-ICOs. As expected, ICOs in both EM and DM are enriched for similar motif sets including but not limited to CTCF, BACH2, ELF3, JUN, HNF4A, and FOXA1, some of which were also enriched in PHH open chromatin loci (Table S2, and Figure 2B). We next applied Gimme maelstrom to analyze differentially enriched motifs on the unique peaks from PHHs, EM- and DM-ICOs. In agreement with our previous observation from SCENIC analysis, we detected high enrichment of TF motifs such as CEBPB, RXRA, PPARG, as well as HNF4A and HNF1A in PHHs. Furthermore, motifs of ELF3, EHF, SOX4/SOX17 and PROX1 were again more enriched in ICOs (Figure 3F). Taken together, our data showed that distinct sets of TFs are required for PHHs and *in vitro* ICOs and identified ELF3 as a TF strongly associated with ICOs.

### ELF3 depletion promotes hepatic differentiation of ICOs in vitro

Next, we focused on ELF3, a TF that is highly expressed and active in ICOs as compared to PHHs, and investigated the effect of depletion of this factor during ICO differentiation. We performed *ELF3* knock-down using a combination of two siRNAs in three different ICO lines at Day 5 after differentiation (in DM), evaluating expression of known hepatocyte markers (Figure 4A). Interestingly, we already observed a strong negative correlation between expression of *ELF3* and expression of *ALB*, *CYP3A4* and *GLUL* within the three ICO clones even without *ELF3* knockdown (Figure 4B). In two of the three clones we observed a robust reduction (∼60% knockdown efficiency on average) of *ELF3* levels in the siRNA transfected samples as compared to the non-targeting siRNA control at differentiation Day 8 (day 3 post-transfection) (Figure 4C). Next, we evaluated expression of known hepatocyte markers and observed increased levels of *ALB*, *CYP3A4*, *TTR*, *GC*, and *GLUL* upon *ELF3* knockdown in both ICO lines tested (Figure 4B, indicated in square and tri-angle). Overall, our data indicates that lower *ELF3* levels associates with more mature hepatic-like features in ICOs under differentiation conditions, suggesting that ELF3 may function as a barrier of hepatic differentiation of ICOs *in vitro*.

**Figure 4.**
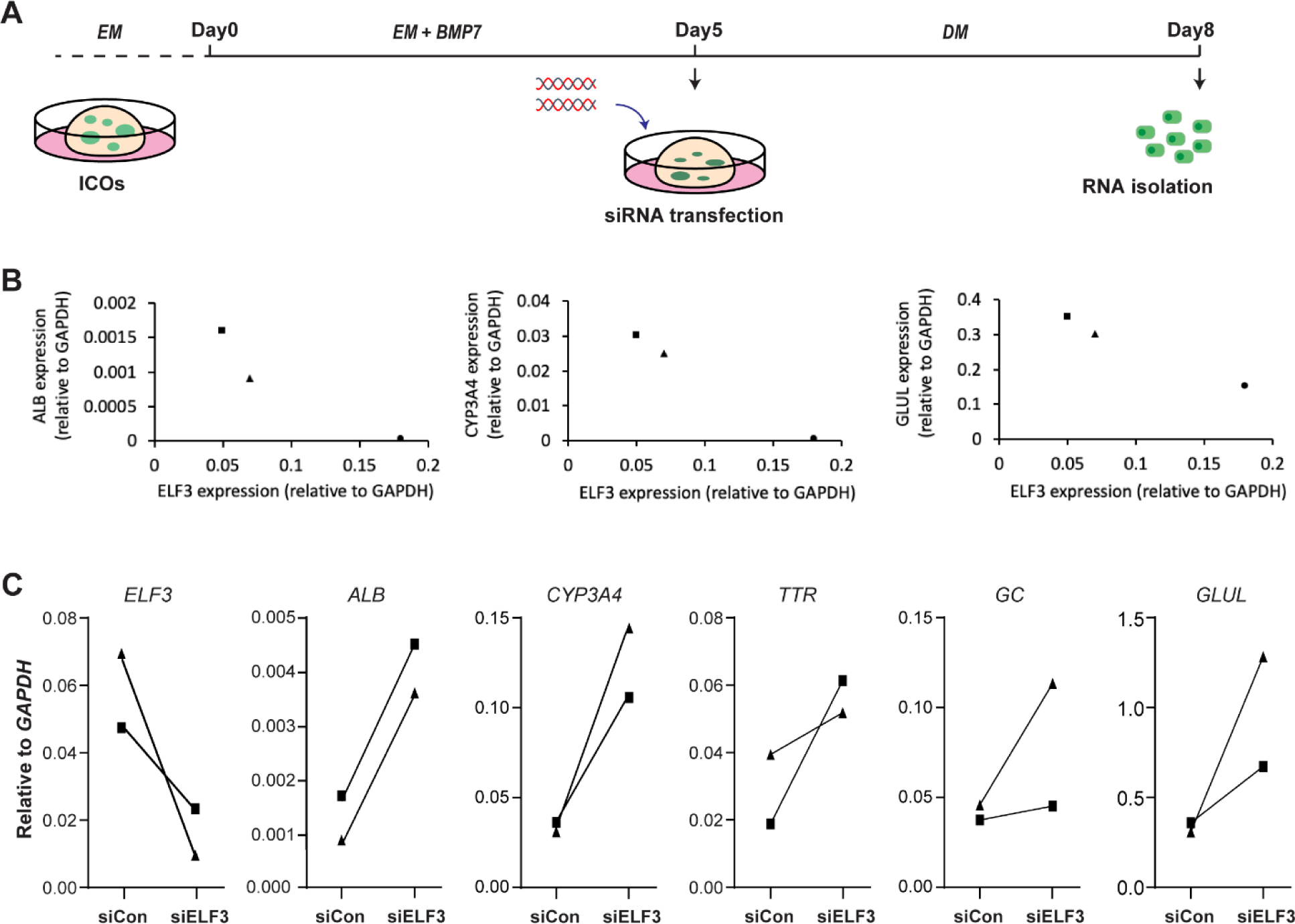
*ELF3* depletion promotes hepatic differentiation in ICOs *in vitro*. **(A)** Experimental setup of siRNA transfection during hepatic differentiation of ICOs. ICOs were maintained in EM. Next, EM-ICOs were first cultured in EM plus BMP7 for 5 days. On day 5 during the differentiation, ICOs were transfected with siRNA (siCon (scrambled siRNAs) and siELF3) and switched to DM medium until day 8 (day 3 post transfection). Cells were harvest for RNA isolation for qPCR. **(B)** RT-qPCR result comparing expression of *ELF3* versus expression of hepatocyte markers *ALB*, *CYP3A4, GLUL* in three different ICO clones after differentiation. **(C)** RT-qPCR result showing the expression levels of *ELF3* as well as hepatocyte markers *ALB*, *CYP3A4*, *TTR*, *GC*, and *GLUL* in *ELF3*-targeting or non-targeting control siRNA transfected DM-ICOs. For panel **B** and **C**: *GAPDH* was used as the house keeping gene.

## Discussion

In this study, we applied an optimized liver tissue dissociation protocol to exclusively isolate primary hepatocytes from human liver. Current available scRNA-seq datasets were mainly obtained from the whole liver to capture all cell types in the tissue (15,16). However, this method usually leads to a limited and inconsistent proportion of PHHs from each donor, causing batch effects in the downstream computational analysis. We modified the two-step perfusion protocol to generate a pure hepatocyte population (98%) from liver tissue. Using scRNA-seq, we confirmed the known zonal heterogeneity within the isolated hepatocytes. Considering that some liver diseases as well as infection have shown zonal preference (32–35), we expect that enriched PHHs (potentially further sorted based on zonal cell surface markers, for which SLCO1B3 could be a candidate) would serve as a better and more precise resource for pericentral liver disease modeling. Laser capture microdissection (LCM) has been used to isolate human hepatocytes from different zones for comprehensive comparison of transcriptome and DNA methylome genome-wide (20). However, LCM extracts both hepatocyte and non-parenchymal cells, and cryosectioning of the cells limits the usage for a broader downstream analysis.

The emergence of liver organoids has offered an alternative *in vitro* long-term culture system to better recapitulate liver tissue in a dish. However, organoid derived hepatic-like cells are functionally limited compared to human hepatocytes, and the underlying biological mechanisms to promote differentiation remain unclear. Therefore, in this study we investigated transcriptomic and epigenomic diversity between PHHs and ICOs. Integrative analysis of scRNA-seq, ATAC-seq and ChIP-seq datasets revealed that besides in liver regeneration, TFs in AP-1 family could play a key role in maintaining cellular functionality of mature PHHs by cooperating with the liver specific TF HNF4A. Although functional investigations are limited due to the lack of a proper model in humans, evidence from other studies could further support our finding. For example, a defect in hepatogenesis has been described in *c-Jun* knockout mice, and *c-Jun* null mouse embryonic stem cells failed to differentiate to hepatocyte in adult livers of chimeric mice (41,42). In addition, a dysregulated level of c-Jun is associated with several liver diseases including hepatitis B virus (HBV) infection, non-alcoholic fatty liver disease (NAFLD), and tumorigenesis (43,44). These results all suggest that proper expression of c-JUN/AP-1 is essential in PHHs. Interestingly, comparative analysis between PHHs and ICOs displayed high expression and activity of AP-1 TFs in both cell types. Motif analysis on unique open chromatin regions showed high enrichment of AP-1 binding motif in ICOs compared to PHHs. AP-1 family contains TFs encoded from the *JUN*, *FOS*, and *ATF* gene families, forming homo/heterodimers and interacting with other DNA binding proteins (45). Our analysis showed that AP-1 TFs cooperate with HNF4A in PHHs. Moreover, cooperarion between AP-1 and ELF3, the key TF in ICOs, on transcription regulation has been described before in other cell types (46–49). Therefore, we reason that different interacting factors could explain the distinct binding profiles of AP-1 TFs in PHHs and ICOs. Alternatively, TF binding is cell-type specific, which is driven by the diverse chromatin states in different cells. The AP-1 motifs enriched in ICO-specific open chromatin regions indicates that TFs in the AP-1 family could have variable regulatory targets and function differently as compared in PHHs. Indeed, JUN and JUNB have been associated with Wnt signaling pathway (50–52), which is highly active in ICOs. Therefore, we hypothesize that AP-1 TFs may function in supporting cell survival and proliferation because of active Wnt pathway in ICOs.

Motif analysis revealed high enrichment of PROX1 and SOX4/SOX17 motifs in ICOs, consistent with their important roles of SOX factors in controlling development of bile duct and cholangiocyte differentiation (53,54). However, the function of PROX1 in cholangiocytes or ICOs are less studied. Previous work has proposed PROX1 as an early marker for the developing liver required for the migration and differentiation of hepatoblasts (55). Liver specific inactivation of Prox1 leads to hepatic injury with very defective hepatocyte morphogenesis (56,57). Since PROX1 is highly expressed in both hepatocytes and ICOs (human liver atlas data, http://human-liver-cell-atlas.ie-freiburg.mpg.de/)(15) with potentially different occupancy preferences, our study has indicated a diverse regulatory function of PROX1 in PHHs and ICOs that requires further functional validation.

Transcription factor ELF3 is highly expressed in biliary epithelial cells (16,58). As reported, our gene regulatory comparison analysis between PHHs and ICOs shows ELF3 to be exclusively active in ICOs, which represent cholangiocyte-like features. Our functional study further reveals upregulation of hepatocyte markers upon *ELF3* depletion in ICOs, suggesting that ELF3 may function as a barrier in hepatocyte differentiation. To date, several studies have reported a role of ELF3 in regulating mesenchymal-to-epithelial/epithelial-to-mesenchymal transitions (MET/EMT), key biological processes that occurs in many physiological events such as development, embryogenesis and cell reprogramming (59–61). In the liver, MET/EMT dynamics are associated with liver progenitor cell identity, hepatocyte differentiation and dedifferentiation upon *in vitro* culture (62–64). Thus, the loss of ELF3 may affect a set of hepatocyte gene expression through MET/EMT processes. RNA-seq analysis using KD of *ELF3* during ICO differentiations, could further elucidate the mode of actions of ELF3 in ICOs. Furthermore, introducing TFs necessary for PHHs, such as MLX and PPARGC1A, into ICOs would offer an alternative strategy for the improvement of functional hepatocyte generation from ICOs, which has been successfully described before in other *in vitro* models (65). In conclusion, our study described the heterogeneity of gene regulatory mechanisms between PHHs and ICOs. These findings provide new biological insights in PHHs as well as ICOs, which will benefit the development of functional human hepatocyte models *in vitro*.

## Acknowledgements

We are thankful to Rob Woestenenk for technical support in FACS sorting experiments. We are grateful to Marijke Baltissen and Lieke Lamers for technical support in next generation sequencing. We thank Sybren Rinzema for sequencing data management. We acknowledge the ENCODE Consortium and the ENCODE production laboratory(s) generating the particular datasets that we included.

## Supplementary Files

Supplementary Table S1: scRNA Differentially expressed genes in each cluster (excel table)

**Supplementary Table S2.**

| *Motifs enriched in EM-ICOs and DM-ICOs* |  |  |  |  |  |
| --- | --- | --- | --- | --- | --- |
| **Top10 Motifs enriched in EM-ICOs** |  |  |  |  |  |
| Motifs | Factors | %input | %background | -log<br>(p-value) | ROC<br>AUC |
| GM.5.0.C2H2_ZF.0024 | CTCF | 11 | <1 | inf | 0.58 |
| GM.5.0.bZIP.0035 | BACH2 | 10 | <1 | 285.42 | 0.62 |
| GM.5.0.bZIP.0013 | JUN | 10 | <1 | 284.06 | 0.62 |
| GM.5.0.bZIP.0058 | NFE2L2 | 4 | <1 | 84.87 | 0.59 |
| GM.5.0.Nuclear_receptor.0028 | HNF4A | 4 | <1 | 74.36 | 0.53 |
| GM.5.0.Ets.0026 | ELF3 | 3 | <1 | 50.05 | 0.59 |
| GM.5.0.CBF_NF-Y.0002 | NFYA | 3 | <1 | 48.22 | 0.55 |
| GM.5.0.Forkhead.0001 | FOXI1 | 3 | <1 | 43.75 | 0.54 |
| GM.5.0.Forkhead.0008 | FOXA1 | 3 | <1 | 42.29 | 0.57 |
| GM.5.0.Unknown.0004 | NF1 | 3 | <1 | 40.11 | 0.49 |
| **Top10 Motifs enriched in DM-ICOs** |  |  |  |  |  |
| Motifs | Factors | %input | %background | -log<br>(p-value) | ROC<br>AUC |
| GM.5.0.C2H2_ZF.0024 | CTCF | 11 | <1 | inf | 0.58 |
| GM.5.0.bZIP.0013 | JUN | 9 | <1 | 250.29 | 0.59 |
| GM.5.0.bZIP.0035 | BACH2 | 7 | <1 | 178.6 | 0.59 |
| GM.5.0.Ets.0026 | ELF3 | 3 | <1 | 57.15 | 0.61 |
| GM.5.0.Unknown.0004 | NF1 | 3 | <1 | 56.99 | 0.51 |
| GM.5.0.Nuclear_receptor.0056 | HNF4A | 3 | <1 | 55.65 | 0.53 |
| GM.5.0.Forkhead.0008 | FOXA1 | 3 | <1 | 39.85 | 0.57 |
| GM.5.0.C2H2_ZF.0288 | SP6 | 2 | <1 | 29.3 | 0.55 |
| GM.5.0.CBF_NF-Y.0002 | NFYA | 2 | <1 | 27.87 | 0.55 |
| GM.5.0.C2H2_ZF.0314 | KLF2 | 2 | <1 | 27.02 | 0.55 |

**Supplementary Table S3.**

| *Patient information* |  |  |  |  |  |
| --- | --- | --- | --- | --- | --- |
| Patient ID | Date of operation | Age | Sex | usage in this manuscript | Type of surgery |
| PHH1 | 2020/9/30 | 48 | Male | scRNA-seq | Wedge resection<br>segment V |
| PHH2 | 2020/10/1 | 72 | Female | scRNA-seq | Wedge resection<br>segment VI |
| PHH3 | 2020/10/14 | 76 | Male | scRNA-seq | Segment VI/VII/VIII<br>and wedge III resection |
| PHH4 | 2021/3/31 | 74 | Male | bulk ATAC-seq | Segment VII resection |
| PHH5 | 2021/4/14 | 38 | Male | bulk ATAC-seq | Wedge resection<br>Segment VIII |

**Supplementary Table S4.**
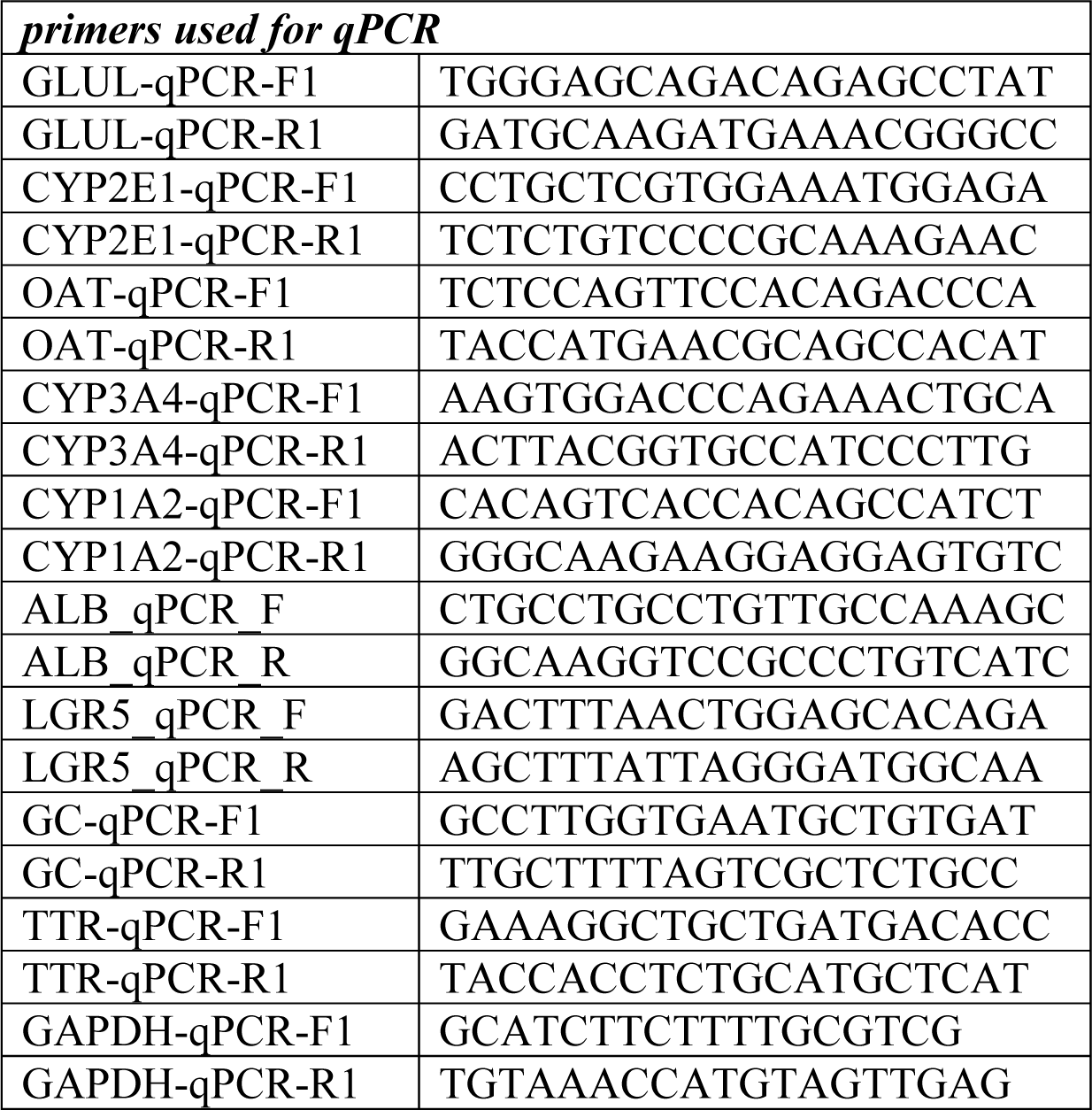

**Supplementary Fig 1.**
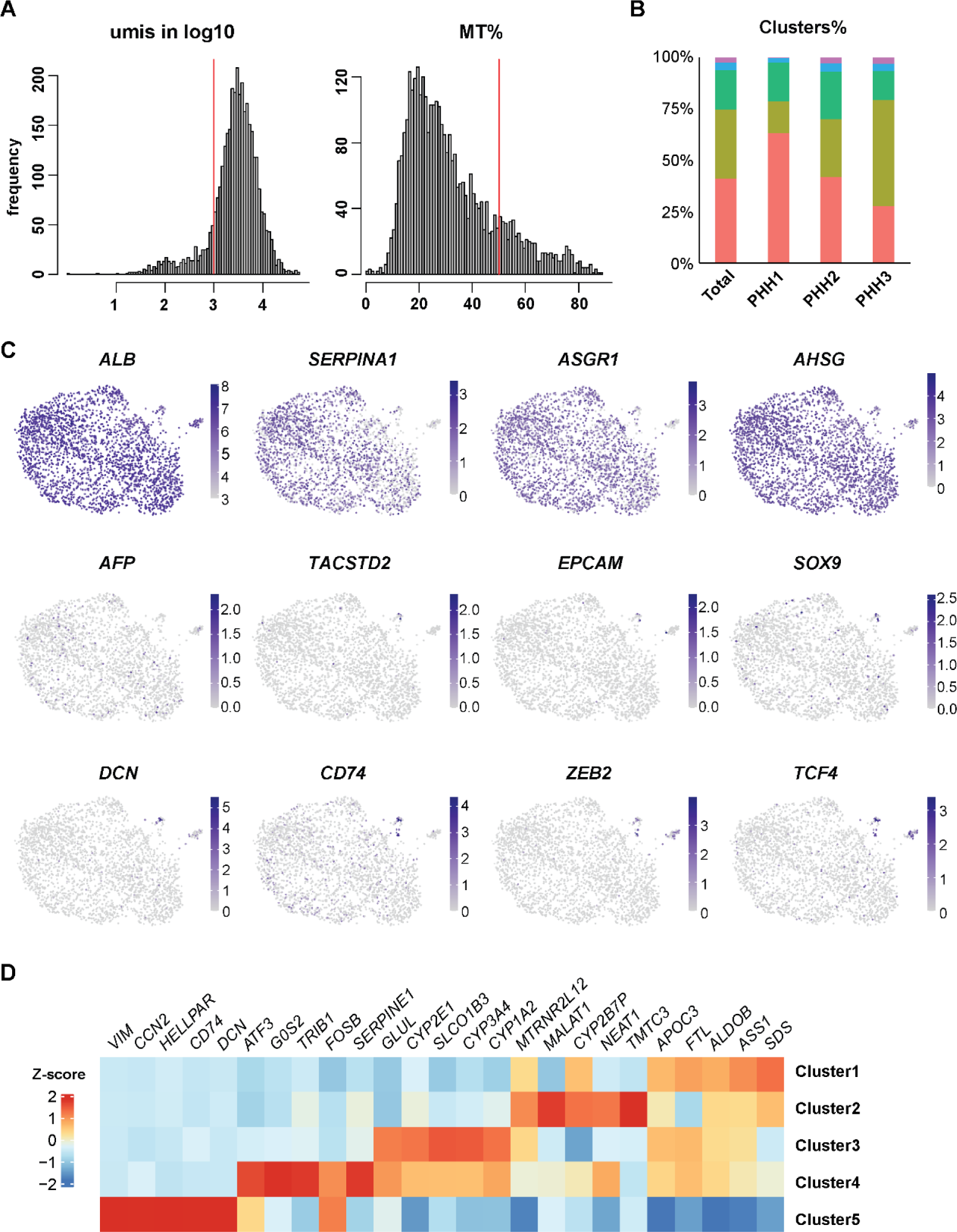
scRNA-seq analysis characterizing liver cells isolated using an optimized two-step perfusion protocol shows nearly pure hepatocyte population. (**A**) Histograms showing the numbers of UMIs (in log10 format) in each cell on the left, and percentage of mitochondrial gene expression per cells on the right. Only the cells following the criteria (1000 < UMIs, MT% < 50%) are used for the downstream analysis. (**B**) Bar plot showing the fraction of cells contributed to each cluster by donors after batch correction. (**C**) UMAP plots of expression of mature hepatocyte markers (*ALB*, *SERPINA1*, *ASGR1* and *AHSG*), liver progenitor markers (*AFP* and *TACSTD2*), cholangiocyte markers (*EPCAM* and *SOX9*), and immune or epithelial cell types associated genes (*DCN*, *CD74*, *ZEB2*, and *TCF4*). Data was normalized to sequencing depth and shown in log2 format. (**D**) Heatmap showing the expression level of top 5 enriched genes from each cluster. Data is shown in Z-score format.

**Supplementary Fig 2.**
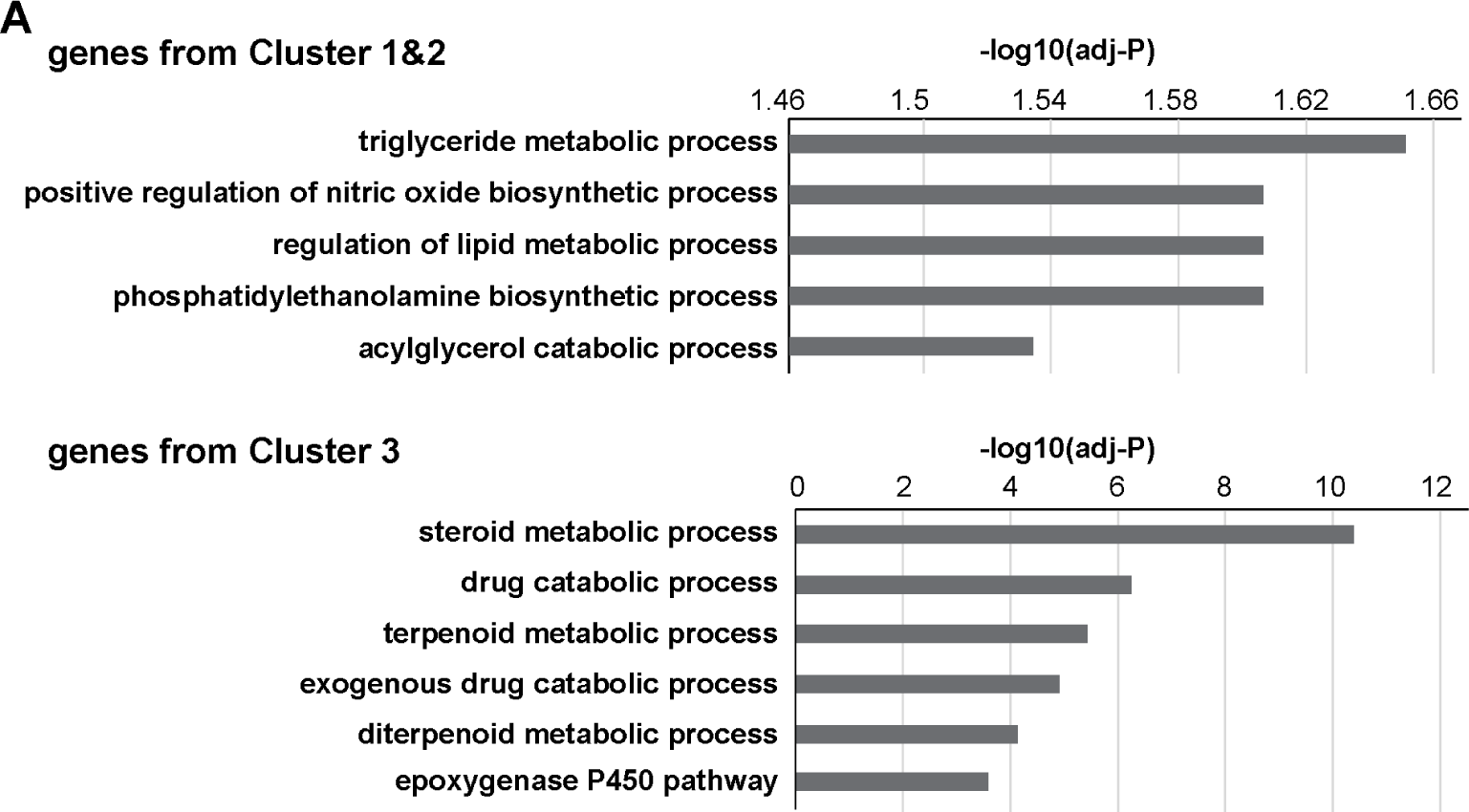
PHH clusters are enriched for typical hepatic processes. (**A**) GO analysis (biological process) for the typical gene sets from cluster 1, 2 and 3. Pathways were listed according to −log10(adj-P value).

**Supplementary Fig 3.**
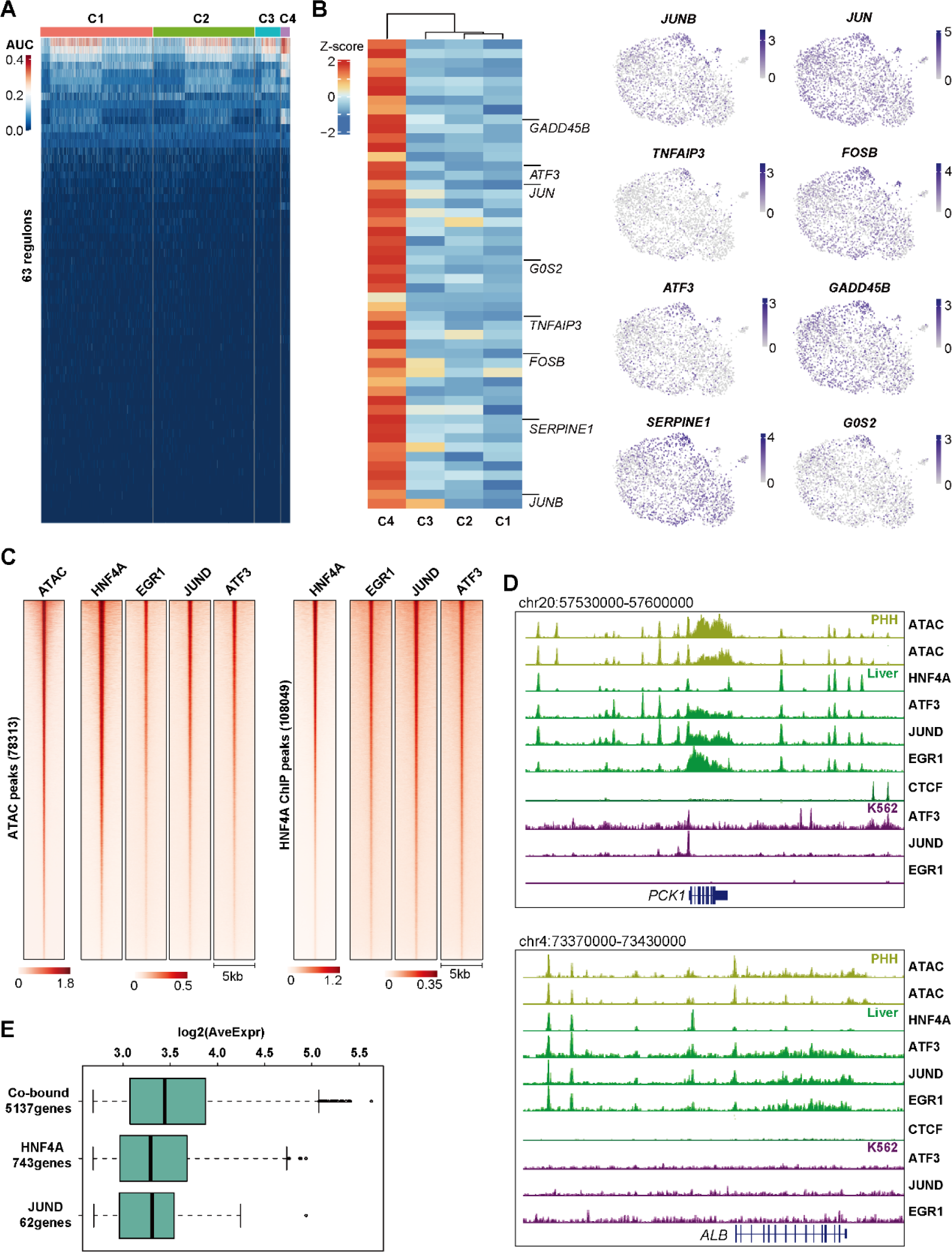
Integrative analysis shows the crosstalk between AP-1 TFs and HNF4A in PHHs. (**A**) Heatmap of the AUC score of all 63 regulons analyzed by SCENIC, showing little difference between clusters. Score was shown in each single cells and cells were grouped in Seurat clusters. (B) Heatmap showing the expression of Cluster4 specific genes in z-score format. Genes associated to hepatocytes in the priming phase are highlighted on the right side (on the left) and UMAP plots displaying the expression levels of *JUNB*, *JUN*, *TNAIP3*, *FOSB*, *ATF3*, *GADD45B*, *SERPINE1* and *G0S2* (on the right). Data was normalized to sequencing depth and shown in log2 format. (**C**) Heatmap showing the enrichment of ATAC signal as well as HNF4A, EGR1, JUND, and ATF3 occupancies in 78313 ATAC peak regions (summit point ±2.5kb) in PHHs (on the left), and the enrichment of HNF4A, EGR1, JUND, and ATF3 occupancies in 108049 HNF4A binding sites (summit point ±2.5kb) in liver (on the right). Data was shown as CPM. (**D**) Genome browser screenshot showing the co-binding of HNF4A, ATF3, JUND, and EGR1 at *PCK1* and *ALB* genomic loci in liver, which overlaps with ATAC signal in PHHs. This co-localization was not observed from either CTCF in liver, or ATF3, JUND, EGR1 in K562 lymphoblast cell line. (**E**) Boxplot showing the average expression level of genes that are bound by both JUND and HNF4A (co-bound), HNF4A only and JUND only. Genes targeted by HNF4A, JUND or both proteins were selected according the following criteria, expression level higher than 100 umis from all summed single cells and the binding sites detected within TSS±5kb of the genes. Data was shown as log2 (total UMIs/ total number of cells in scRNA-seq *1000)

**Supplementary Fig 4.**
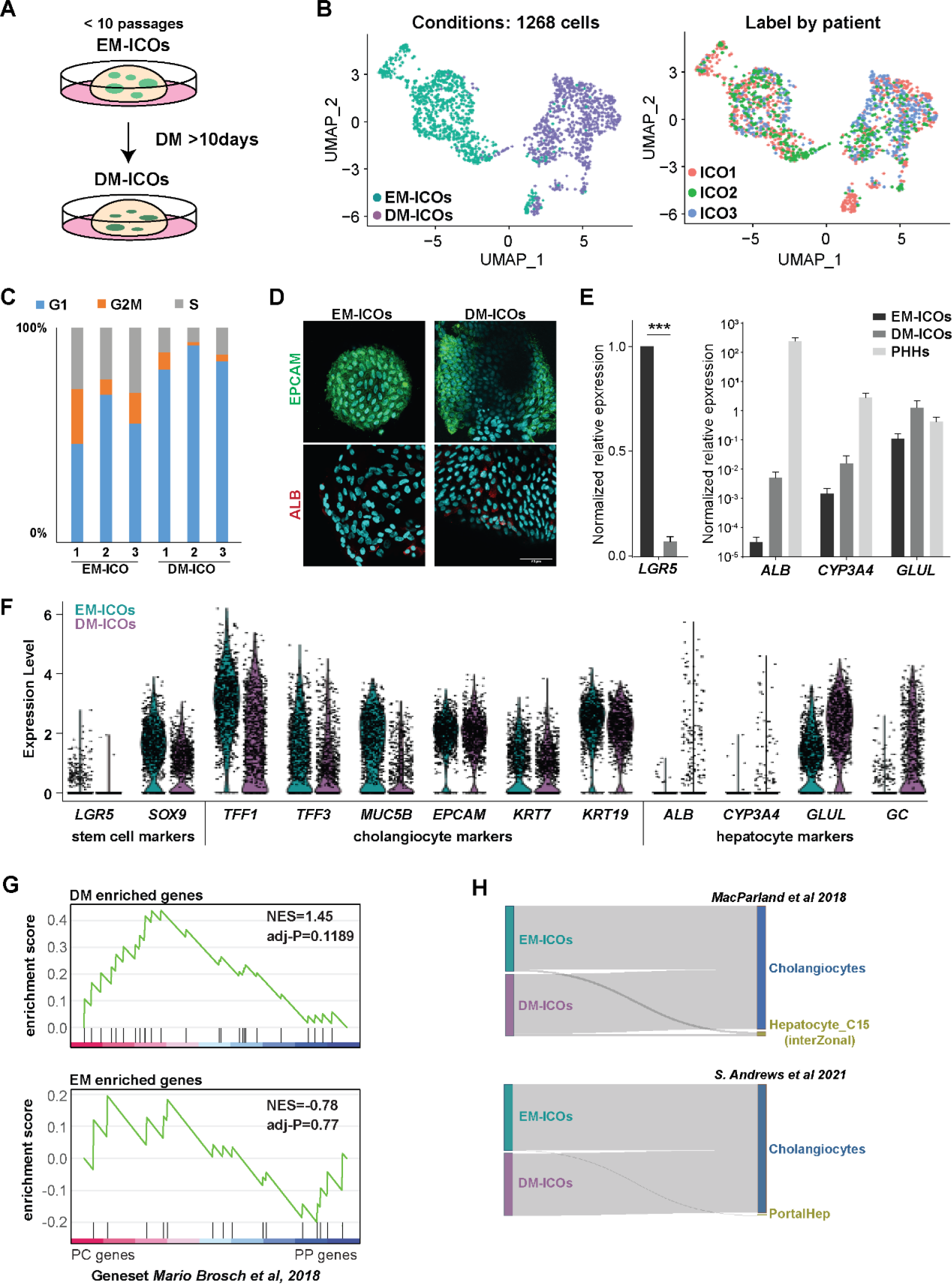
DM-ICOs display hepatocyte features compared to EM-ICOs. (**A**) Schematic overview of the ICO culture. ICOs were maintained in EM. For hepatic differentiation, ICOs were cultured in DM for more than 10 days. All the experiments were performed using the ICO lines within 10 passages. (**B**) UMAP plots of single-cell transcriptome labeled by culture condition (left) or donors (right). Cells from different donors are separated mainly by the culture condition. (**C**) Bar plot showing the distribution of cells from EM-ICOs and DM-ICOs in different cell-cycle phases (G1, S, G2M). Cells were grouped based on the donors. (**D**) Immunostaining of EPCAM and ALB in EM-ICOs and DM-ICOs. Hoecht staining for the nuclei was used as a control. (**E**) qPCR results showing the expression levels of *LGR5*, *ALB*, *CYP3A4*, and *GLUL* in PHHs, EM-ICOs, and DM-ICOs. *GAPDH* was used as the house keeping gene. Expression was normalized to EM-ICO samples. Data was generated with 3 biological replicates. Mean ± SEM. Student t-test \*\*\**P<0.001*. (**F**) Violin plots showing the expression of selected genes such as WNT target gene *LGR5*, progenitor/cholangiocyte marker *SOX9*, mature cholangiocyte markers (*TFF1*, *TFF3*, *MUC5B*, *EPCAM*, *KRT7*, and *KRT19*), and mature hepatocyte markers (*ALB*, *CYP3A4*, *GLUL*, and *GC*). Cells were grouped based on culture conditions. (**G**) GSEA of DE genes between EM-ICOs and DM-ICOs compared to a zonal hepatic gene set obtained from a published study, showing that DM or EM genes were not enriched to any zonal expression profile. (**H**) Sankey plots showing the predicted cell types of ICOs in this study by Random Forest classifier based on transcriptomes of liver cells (hepatocytes and cholangiocytes) from two independent studies.

**Supplementary Fig 5.**
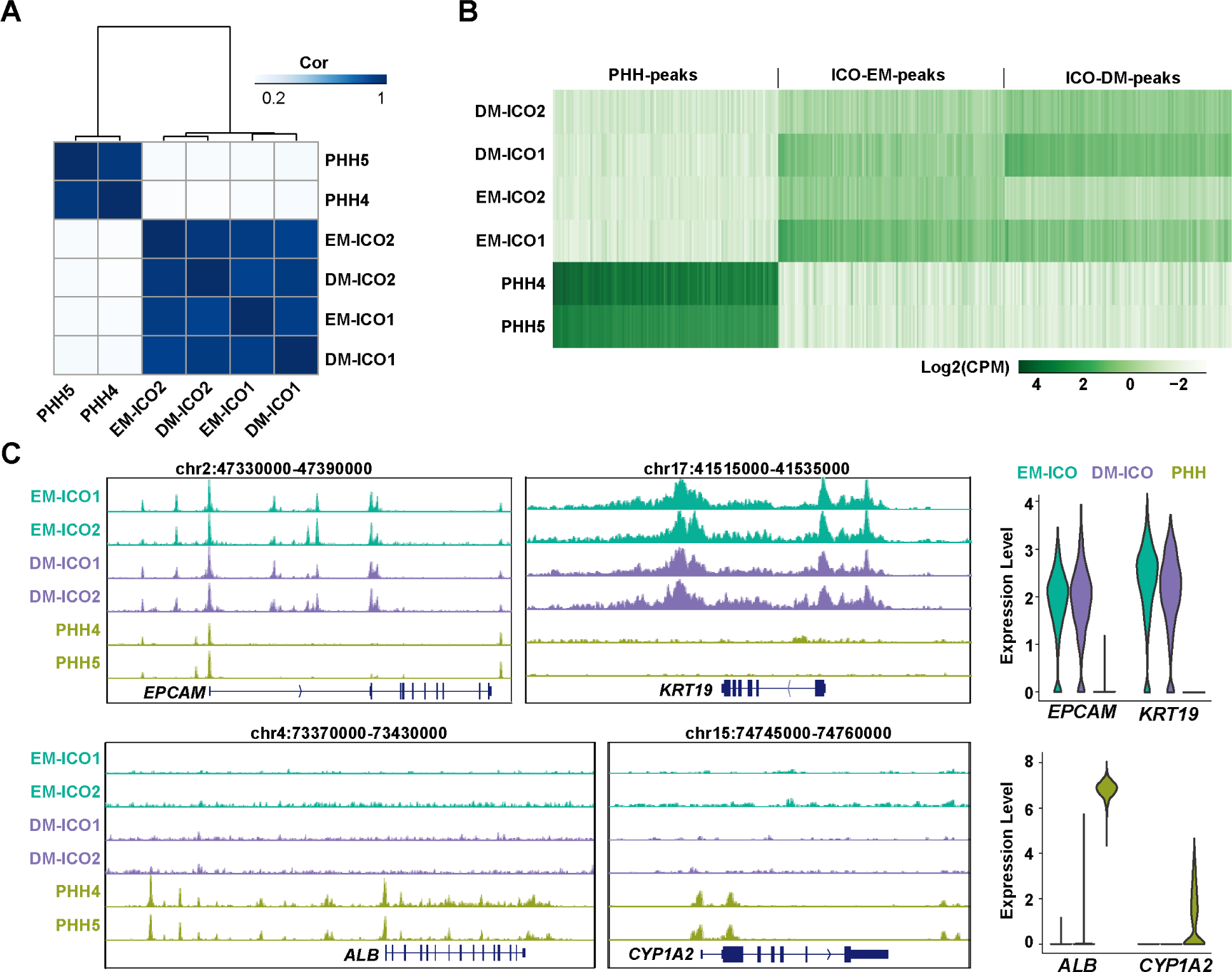
Comparative analysis of PHHs and ICOs shows distinct chromatin profiles, and identified ELF3 as a barrier of hepatic differentiation in ICOs. (**A**) heatmap of spearman correlation matrix of normalized open chromatin intensities in PHHs, EM-ICOs, and DM-ICOs identified by ATAC-seq. (**B**) Heatmap showing the difference of chromatin accessibility (top 2000 ATAC peaks of each sample) between PHHs and ICOs in EM and DM conditions. Data was shown in Log2(CPM). (**C**) Genome browser screenshot showing the chromatin state at *EPCAM*, *KRT19*, *ALB*, and *CYP1A2* genomic loci on the left, and the violin plots showing the corresponding expression levels in PHHs and ICOs on the right.

